# Intersubject information mapping: revealing canonical representations of complex natural stimuli

**DOI:** 10.1101/016436

**Authors:** Nikolaus Kriegeskorte

**Affiliations:** Medical Research Council, Cognition and Brain Sciences Unit, 15 Chaucer Road, Cambridge, CB2 7EF, UK, www.mrc-cbu.cam.ac.uk/people/nikolaus.kriegeskorte

## Abstract

Real-world time-continuous stimuli such as video promise greater naturalism for studies of brain function. However, modeling the stimulus variation is challenging and introduces a bias in favor of particular descriptive dimensions. Alternatively, we can look for brain regions whose signal is correlated between subjects, essentially using one subject to model another. Intersubject correlation mapping (ICM) allows us to find brain regions driven in a canonical manner across subjects by a complex natural stimulus. However, it requires a direct voxel-to-voxel match between the spatiotemporal activity patterns and is thus only sensitive to common activations sufficiently extended to match up in Talairach space (or in an alternative, e.g. cortical-surface-based, common brain space). Here we introduce the more general approach of intersubject information mapping (IIM). For each brain region, IIM determines how much information is shared between the subjects' local spatiotemporal activity patterns. We estimate the intersubject mutual information using canonical correlation analysis applied to voxels within a spherical searchlight centered on each voxel in turn. The intersubject information estimate is invariant to linear transforms including spatial rearrangement of the voxels within the searchlight. This invariance to local encoding will be crucial in exploring fine-grained brain representations, which cannot be matched up in a common space and, more fundamentally, might be unique to each individual – like fingerprints. IIM yields a continuous brain map, which reflects intersubject information in fine-grained patterns. Performed on data from functional magnetic resonance imaging (fMRI) of subjects viewing the same television show, IIM and ICM both highlighted sensory representations, including primary visual and auditory cortices. However, IIM revealed additional regions in higher association cortices, namely temporal pole and orbitofrontal cortex. These regions appear to encode the same information across subjects in their fine-grained patterns, although their spatial-average activation was not significantly correlated between subjects.

## 1 Introduction

The brain is highly adapted to the natural world. To understand its mechanisms we need to study its behavior under natural conditions. One important step in that direction is the use of time-continuous natural stimuli. For example, videos of the real world provide useful stimuli, because they are more natural in their spatiotemporal audio-visual structure than the stimuli conventionally used in neuroscientific experiments.

In order to learn whether response patterns in a given brain region reflect the stimulus, we can build a model of the stimulus variations along time. However, complex natural stimuli vary on an infinite number of dimensions. We could model low-level image statistics or high-level semantic information, such as the presence of particular characters in the scene. At either level, there is an unlimited number of dimensions of which we must choose a small number, because we are limited by the temporal complexity of our data set. Whatever choice we make will strongly bias our results.

An alternative approach to this problem is to "model" a given subject's brain responses by the responses of other subjects. When two people are watching the same television show in isolation, any mutual information between their brain responses must be caused by the common stimulus. The only assumption this inference requires is that the subjects are not telepathically connected (Fig. 1a, b). We can therefore determine if a brain region reflects some aspect of a common stimulus stream (e.g. television) by testing for mutual information between the two brains. (The brain responses, of course, need not be measured simultaneously in different subjects, as long as analysis is time-locked to the stimulus stream.)

**Fig. 1.**
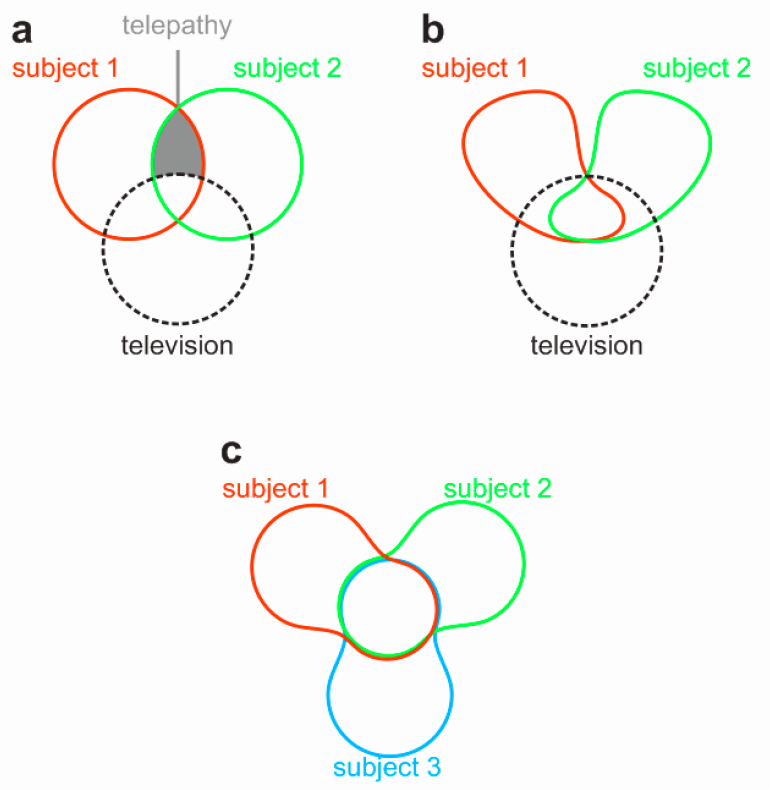
Venn diagrams of mutual information between subject brains and a common complex input. The areas of the shapes represent the information in a stimulus stream or in a subject's brain. Overlap represents mutual information. (**a**) When two subjects (red, green) perceive the same stimulus stream (television), their brains share information time-locked to the stimulus stream. If the subjects are isolated from each other, any mutual information between their brains (red-green overlap) must reflect either the common input (television), or a telepathic connection (gray). (**b**) Assuming that the subjects are not telepathically connected, any intersubject information (red-green overlap) must reflect the common input (television). (**c**) Given only brain activity time-locked to a common stimulus stream for a group of subjects, but no description of the stimulus, we can infer whether a given brain region is driven by some aspect of the stimulus stream by detecting intersubject information.

Hasson et al. (2004) demonstrated that different subjects viewing the same movie exhibit correlated brain activity in visual, auditory, and association cortices. This approach is free of the bias associated with the specification of a design matrix that describes the complex stimulus by a limited set of descriptive dimensions. However, the method of intersubject correlation mapping (ICM) used by Hasson et al. (2004) requires a direct voxel-to-voxel match between the spatiotemporal activity patterns. It is thus only sensitive to common activations sufficiently extended in the brain to match up in Talairach space (or an alternative, e.g. cortical-surface-based, common brain space). The method exploits fluctuations of regional-average activation, but cannot reveal mutual information in fine-grained activity patterns within each functional region.

Here we introduce the more general approach of intersubject information mapping (IIM). For each brain region, IIM determines how much information is shared between the subjects' local spatiotemporal activity patterns. To assess intersubject information, we perform canonical correlation analysis, which measures the overall linear dependency between two sets of variables (regional responses in subject 1 and 2), and has a simple relationship to mutual information. In order to make a continuous brain map highlighting regions that reflect the stimulus stream, we assess the intersubject mutual information for voxels within a spherical searchlight centered on each voxel in turn. This yields a continuous map of intersubject information, which is sensitive to fine-grained representations and yet robust to the imprecision of Talairach spatial correspondence.

At a given location, the information estimate does not change if voxels within one subject's spherical searchlight are spatially rearranged, inverted in sign, or otherwise linearly transformed without loss of information. IIM therefore does not require that subjects share the same fine-grained activity pattern, as long as common information is encoded in a given region. These invariances will be crucial in exploring fine-grained representations, which cannot be matched up in a common space and, more fundamentally, may be unique to each individual – like fingerprints. IIM addresses a challenge that will gain in importance as high-field fMRI, electrode-array recordings, and other techniques of high-resolution multichannel brain-activity measurement become more widely used.

We demonstrate IIM by applying it to fMRI data of three subjects viewing the same segments of a TV series. The technique detected a widely distributed network of brain regions including primary visual and auditory regions, higher-level perceptual regions, and frontal and parietal cortices. Importantly, IIM revealed additional regions not detected by ICM.

## 2 Methods

### 2.1 Measuring the information shared between two regions

A given brain region could encode the same information in subject 1 and 2 by means of a different code (Figs. 2, 3). At the simplest level, the units might encode the same set of representational dimensions in both subjects, but reside in different locations within the region. More generally, the codes may be related by some linear or nonlinear transform. There is an infinite number of possible codes for the same information. The goal of intersubject information mapping is to abstract from this variability and reveal the mutual information.

**Fig. 2.**
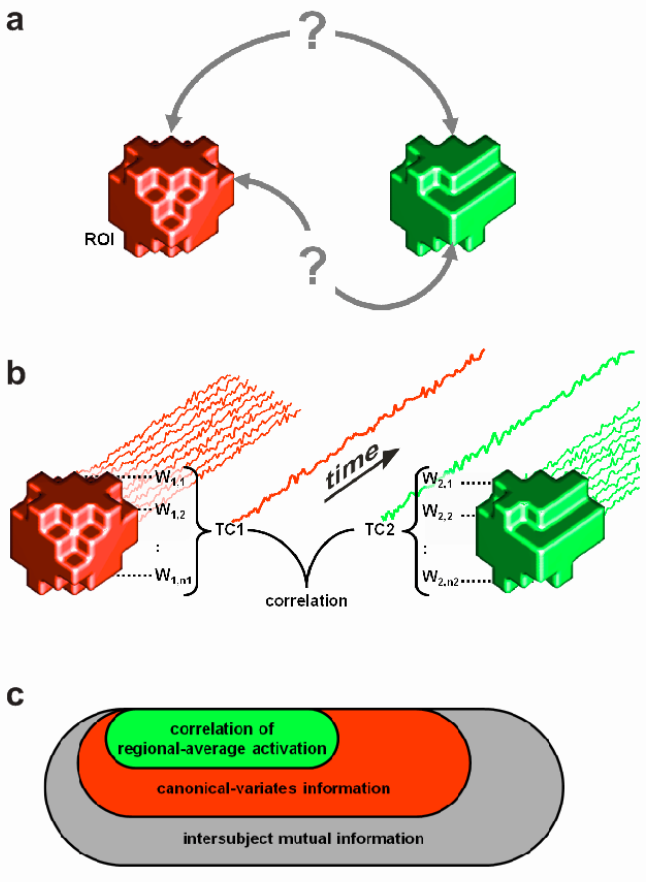
Measuring shared information in corresponding functional regions of two subjects with canonical correlation analysis. (**a**) Given spatiotemporal brain activity patterns in functionally corresponding brain regions in two subjects (red, green), it is difficult to relate the activity patterns because the fine-grained spatial layout of the neuronal patterns might not match up in a common brain space and, more fundamentally, the spatial layout might be unique to each subject like a finger print. (**b**) Canonical correlation analysis finds weights for each response channel (e.g. voxel) in each subject, such that the correlation between the weighted-average time courses is maximized. This "first canonical correlation" provides a linear first approximation to the information shared between the regions. (**c**) Canonical-variates information (red) is a superset of regional-average correlation information (green) and a subset of the total mutual information (gray) between the corresponding brain regions in the two subjects.

**Fig. 3.**
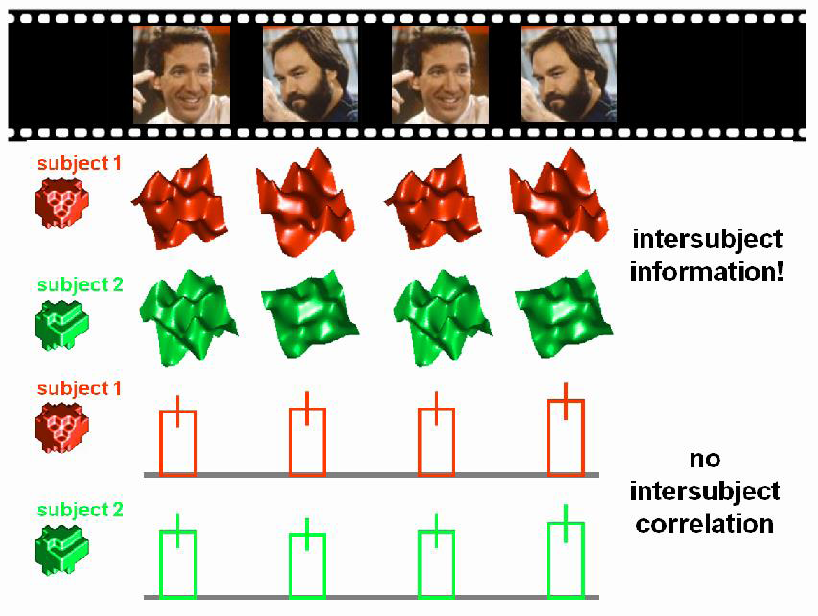
Intersubject information does not imply intersubject correlation. A given brain region in two subjects (red, green) can encode the same information about a common stimulus stream (television clip at the top), even if the spatial brain-activity patterns encoding the information differ between the subjects, and even if the regional-average activation is constant across time or uncorrelated between the subjects.

Estimating mutual information without strong restrictive assumptions requires many data points in high-dimensional scenarios. In fMRI, the number of time points is usually small (in the hundreds), whereas the number of voxels (i.e. the number of dimensions) is large (tens of thousands). In the present scenario, our aim is to assess the mutual information between subjects not for the entire brain at once, but for one region at a time. This reduces the number dimensions to tens or hundreds of voxels. Even for small regions, however, mutual information estimators that make minimal assumptions about the nature of the relationship between the variables (Kraskov et al. 2004) give noisy estimates in typical fMRI scenarios. This is an inevitable consequence of the large space of possible nonlinear relationships between the variables. Here we restrict our sensitivity to mutual information between the subjects to linear relationships. This greatly improves the stability of the estimates.

The linear relationship between two unidimensional variables is measured by the correlation coefficient. The linear relationship between a unidimensional variable and a multidimensional variable is measured by the multiple correlation coefficient. These two scenarios are special cases of canonical correlation analysis (CCA), which measures the linear relationship between two multidimensional variables (Hotelling 1936; Borga 1998; for an application in fMRI activation mapping, see Friman et al. 2001).

CCA jointly optimizes voxel weights *w_1,1_, w_1,2_,... w_1,n1_* (for the *n1* voxels of the region in subject 1) and *w_2,1_, w_2,2_,... w_2,n2_* (for the *n2* voxels of the region in subject 2), so as to maximize to linear correlation coefficient r between the univariate time courses *TC1* and *TC2* that obtain as the weighted sums of the voxel time courses computed for subject 1 and subject 2, respectively (Fig. 2b).

When both variables are multidimensional, as in our scenario here, the shared variance is also multidimensional and not fully captured by a single correlation. Similar to principal components analysis, CCA determines subsequent dimensions of shared variance, each orthogonal to all previous ones and associated with a correlation coefficient that doesn't exceed the previous one.

If the joint probability density of the variables is Gaussian, then the mutual information between regions *R_1_* and *R_2_* (corresponding brain regions in two subjects here) is:

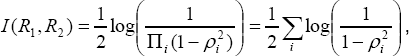

where *ρi* is the *i*-th canonical correlation (Borga 1998). The simple sum in the right-hand formula obtains because the dimensions of shared variance are orthogonal to each other. This estimate of mutual information for the multinormal scenario provides a useful and well-motivated scalar summary of the multiple canonical correlations.

The decomposition of the shared variance into orthogonal components explaining decreasing variance is useful in the present scenario for two reasons: (1) It allows us to control the bias-variance tradeoff by focusing the sensitivity of the method on a restricted number of dimensions. Excluding dimensions explaining little variance will stabilize the estimates. (2) Extracting one or a small number of dimensions of shared variance might allow us to examine the shared-variance time course, relate it back to the stimulus stream, and discover the content of the representation.

For stability of the estimates, the analyses in this paper use only the first canonical correlation. The multinormal estimate of the mutual information grows monotonically with the first canonical correlation, and we use the latter as the test statistic of "intersubject information".

### 2.2 Intersubject correlation and information as complementary measures

In order to directly compare intersubject correlation (i.e. correlation of fluctuations of regional-average activation) and intersubject information (i.e. canonical correlation of fine-grained activity patterns), we compute both statistics within a given region of interest. To compute the intersubject correlation, we first average all voxel time courses within the region in each subject and then compute the correlation of the resulting time courses between subjects.

Intersubject correlation reflects only mutual information encoded in the regional-average activation, and only if it is encoded in the same way (only positive correlations are considered). Intersubject information mapping with CCA is sensitive to any linear relationship between the codes, including intersubject regional-average correlation as a special case, but excluding nonlinear relationships (Fig. 2c).

In order to obtain complementary measures of the regional-average-activation-based intersubject correlation and the fine-grained-pattern-based intersubject information, we remove the activation fluctuations before CCA. To remove activation fluctuations in each subject, we consider each time point in turn and subtract the regional average from each voxel's value at that time point. After removing the activation fluctuations, the regional-average time course consists only of zeros.

### 2.3 Searchlight-based intersubject information mapping

In order to make a brain map of intersubject information, we move a searchlight (Kriegeskorte et al. 2006; Kriegeskorte & Bandettini 2007) through the brains of all subjects in parallel (Fig. 4). The searchlight is placed at corresponding brain locations in all subjects at each step. Here we use a volume-based approach with a voxel sphere as the searchlight and with Talairach space defining the spatial correspondency mapping. Alternatively, a cortical-surface-based approach with a surface-patch searchlight could be used (Oosterhof et al. 2010a, 2010b).

**Fig. 4.**
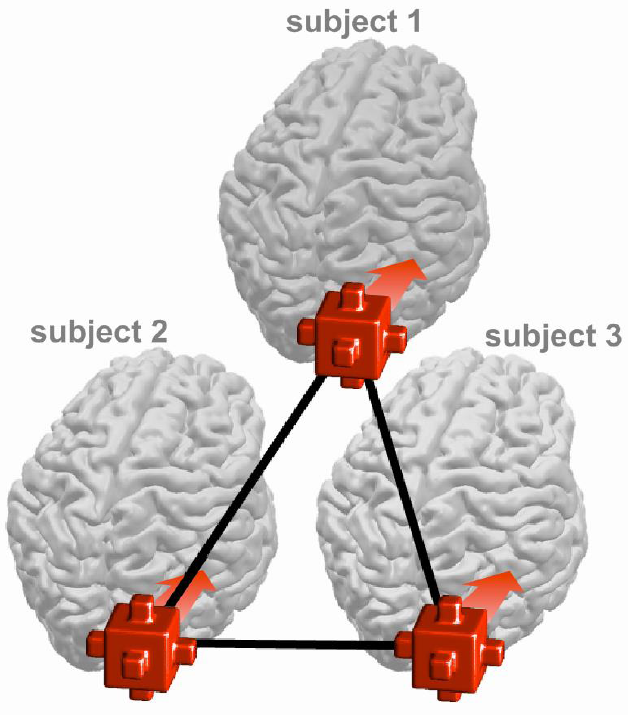
Intersubject information mapping with searchlights in a common space. We can find brain regions encoding common information in multiple subjects by moving spherical searchlights through their brains and analyzing for intersubject information in the local activity patterns at each location. In the simplest implementation, the searchlights are placed at corresponding locations in a common brain space such as Talairach space at each step.

We perform searchlight mapping of (1) intersubject correlation (correlating the time courses obtained by averaging all searchlight voxels within each subject) and (2) complementary intersubject information (estimating the first canonical correlation as the statistic of "intersubject information", after removing the regional-average activation fluctuations). These two maps are computed for each pair of subjects. We then average the maps across all pairs of subjects to obtain one overall intersubject correlation map and one overall intersubject information map.

### 2.4 Statistical inference on intersubject information maps

In CCA, statistical inference can be performed readily if we assume multivariate normality. However, we do not want to rely on multivariate normality for the validity of statistical inference here. Although univariate normality of the errors often approximately holds in voxel-by-voxel linear regression analysis of fMRI data, we have found significant violations of multivariate normality for fMRI regions of interest in previous unpublished analyses (see also Kriegeskorte et al. 2006; Kriegeskorte 2011).

In order to test whether the intersubject information statistic is significant, we therefore simulate the null hypothesis (i.e. no intersubject information) by temporally scrambling the data from one of the two subjects of each subject-pair analysis (details below). The simulation is repeated many times so as to generate a distribution of the intersubject information statistic under the null hypothesis. The p value then obtains as the proportion of simulated-null intersubject information statistics that exceed the actual intersubject information statistic for a given region. We first use searchlight mapping to produce a map of p values, uncorrected for multiple testing. Then we threshold the map so as to control the false discovery rate (Benjamini & Hochberg 1995; Genovese et al. 2002) at 5%. The false discovery rate is the expected proportion of falsely highlighted voxels among all highlighted voxels.

If the goal were to control the familywise error rate (i.e. the probability of highlighting even a single voxel under the null hypothesis) instead of the false-discovery rate, we could use the same temporal scrambling approach to simulate the distribution of searchlight map maxima of the intersubject information statistic under the null hypothesis (Nichols & Holmes 2002; Nichols & Hayasaka 2003). To control the familywise error rate at 5%, we would choose a threshold exceeded by only 5% of the simulated null maps.

In the present analyses the intersubject information statistic is the first canonical correlation. However, the inference method is equally valid for other intersubject information statistics such as the multinormal mutual information estimate described above (which could be computed for the first, the first m, or all canonical correlations to tradeoff bias against variance).

The goal of temporally scrambling the data of one subject is to disrupt the temporal synchrony between the subjects (thus obliterating the mutual information), while preserving the spatiotemporal dependency structure of the data. A simple approach would be to cut the data into a number of temporal blocks and randomly rearrange the temporal sequence of the blocks. Within each block, the spatiotemporal dependencies would be preserved. However, at the borders between blocks, we would introduce discontinuities untypical of a continuous sequence. (If the blocks are large and the borders thus few, this might be negligible.)

Here we use a whitening resampling approach. We fit a first-order autoregression model (AR(1)) to each time series (e.g. Bullmore et al. 1996; Worsley et al. 2002; Friman & Westin 2005). We then randomly permute the whitened residuals (keeping each time point's residual vector across voxels intact to preserve spatial dependencies). Finally we reimpose the temporal dependency of the original time series.

### 2.5 Experiment, fMRI measurements, preprocessing, and analysis

*Experiment*. As part of the “Experience-based cognition” (EBC) project at the University of Pittsburgh, subjects were presented with video clips taken from the TV series Home

Improvement while their brain activity was measured with fMRI. Three 18-minute runs were acquired in each of three subjects. Each subject viewed the same sequences of the TV series (but didn't view any part repeatedly) while being scanned.

*fMRI measurements.* The data were acquired using a 3-Tesla Siemens Allegra scanner and an echo-planar imaging sequence for blood-oxygen-level-dependent fMRI. The time-to-repeat from one volume to the next was 1.75 s. 34 slices were acquired in axial orientation with a field of view of 210 mm width and a 64x64 grid, yielding a voxel size of (3.28125 mm)^2^ with a slice thickness of 3.5 mm and no gap between slices. The time-to-echo was 25 ms and the flip angle was 76 degrees.

*Preprocessing.* The data were preprocessed using the BrainVoyager analysis software. The preprocessing consisted in head-motion correction (by rigid body alignment and resampling of each functional volume), slice-scan-time correction (by linear interpolation in time, bringing the sequentially acquired slices within each volume into register in time), and linear trend removal (applied to each voxel time course separately). The fMRI data were transformed on to Talairach space and resampled into an isotropic (3 mm)^3^ voxel grid in this process. Head-motion parameter time courses (3 rotational and 3 positional parameters) were regressed out of each voxel time course to further reduce residual (e.g. non-rigid) head-motion artifact.

*Analysis.* Intersubject information mapping was performed within a Talairach-space brain mask using 5-mm radius (19 voxel) searchlights placed at corresponding locations in Talairach space in each subject in each step. Analyses used the first canonical correlation between corresponding searchlight locations computed for each pair of subjects and each run, and then averaged across all pairs of subjects and all runs. Inference was performed using the method described in detail above. We repeated the intersubject information mapping 1000 times on simulated null data generated using whitening resampling to temporally scramble the time series. This provided a simulated null distribution of the grand-average first canonical correlations (unbiased by weighting and without normal assumptions) from which we computed a single p map. This p map was thresholded so as to control the false-discovery rate at 5%.

## 3 Results

Fig. 5 shows the intersubject information map (based on the first canonical correlation after subtracting out the searchlight-average activation for each time point in each subject). Regions where information is shared between subjects include early visual areas, lateral occipital complex (LOC), the fusiform face area (FFA), the parahippocampal place area (PPA), the human motion complex, superior parietal regions, intraparietal sulcus, precuneus, superior temporal sulcus, primary and higher auditory cortex, temporal pole, orbitofrontal cortex, superior medial frontal cortex, and right precentral gyrus.

**Fig. 5.**
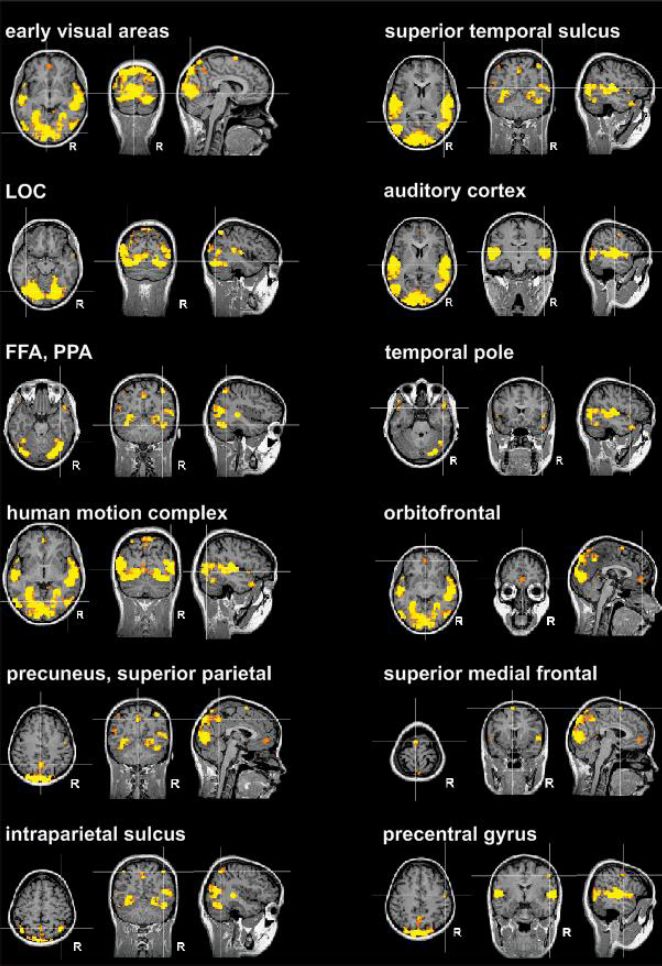
Intersubject information map. In each panel, an axial, a coronal, and a saggital slice intersect in a different cortical region (panel label). In each slice, the positions of the other two slices of the panel are marked by white lines forming crosshairs. The intersubject information statistic is the first canonical correlation between searchlights placed at corresponding locations in Talairach space. The searchlight-average activation has been subtracted out for each time point in each subject. The map is thresholded to control the false discovery rate at 5%.

Fig. 6 compares the intersubject information map to the intersubject correlation map. Many voxels are highlighted in both maps (yellow dots in scatterplot). The regions where these voxels are located include early visual cortex, early auditory cortex, and ventral temporal cortex. The intersubject information map highlights many voxels not highlighted in the intersubject correlation map (red dots in scatterplot). The regions where these voxels are located include temporal pole and orbitofrontal cortex.

**Fig. 6.**
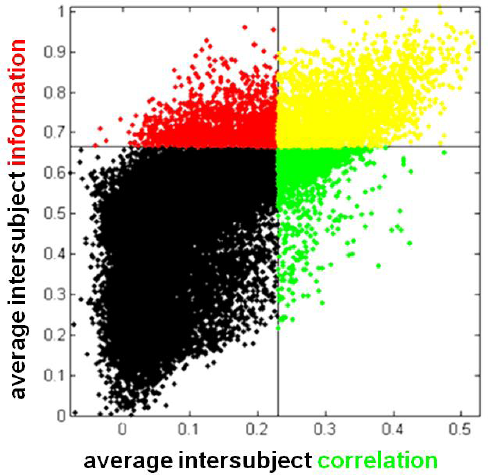
Relationship between intersubject correlation and intersubject information maps. Each dot in the scatterplot corresponds to a voxel in Talairach space. The horizontal position of each dot is determined by the intersubject correlation (searchlight-average activation correlation computed for each pair of subjects and averaged across all subject pairs). The vertical position of each dot is determined by the intersubject information (first canonical correlation between searchlights after searchlight-average activation has been subtracted out for each time point in each subject). The dividing lines correspond to the thresholds controlling the false discovery rate at 5%. Yellow dots correspond to voxels highlighted in both maps, red dots to voxels highlighted only in the intersubject information map, green dots to voxels highlighted only in the intersubject correlation map. Note that the overall shape of the dot distribution approximates a triangle: Low intersubject correlation says little about intersubject information. However, low intersubject information implies low intersubject correlation.

Significant intersubject information is frequently found in regions with very low intersubject correlation. However, the reverse is not true: When the intersubject information is very low, then the intersubject correlation is also low. Despite this tendency, there are some voxels that are highlighted only in the intersubject correlation map. These tend to have intermediate intersubject information.

Overall these results reflect the fact that intersubject information is a more general measure including intersubject correlation as a special case. We subtracted out the searchlight-average activation to make the measures more complementary. However, results suggest that where there is intersubject correlation there also tends to be information beyond covariation of the local-average activation. Voxels highlighted only in the intersubject correlation map (green dots in scatterplot) can result from a combination of (a) regions with intersubject correlation, but no fine-grained pattern information, (b) the noise in the data randomly pushing the correlation statistic, but not the information statistic above the threshold, (c) the fact that intersubject correlation is easier to distinguish from noise as it constitutes a more specific hypothesis.

## 4 Discussion

### 4.1 Common information represented in corresponding brain regions of different subjects

Intersubject information mapping detected significant evidence of intersubject information in a large set of cortical regions, with strong effects not only in primary visual and auditory cortices, but also in higher-level association cortices distributed across all cortical lobes. We focus the discussion on primary sensory regions found in both intersubject correlation and intersubject information mapping, and on the temporal pole and orbitofrontal regions found only in the intersubject information mapping.

Primary visual and auditory areas exhibited both intersubject correlation and intersubject information. These findings are expected because the two areas are known to be directly driven by the audiovisual stimulus stream. The correlation of the regional-average activation likely reflects the overall stimulus energy in each modality (integrated over visual space and auditory frequency bands, respectively). The canonical correlation of the activity patterns likely reflects the representation at each point in time of the current visual and auditory stimulus pattern, which all subjects share.

Temporal pole and orbitofrontal cortex were found in the intersubject information, but not in the intersubject correlation mapping, suggesting that these regions might contain common information across subjects in their more fine-grained patterns of activity, whose spatial structure is likely to be subject-unique and will not match up in Talairach space. Both of these regions integrate complex sensory and endogenous information. Anterior temporal regions are thought to relate object perception and memory, contributing to the recognition of familiar faces and objects (Nakamura & Kubota 1996; Olson et al. 2007; Kriegeskorte et al. 2007). Orbitofrontal cortex is thought to predict behavioral outcome values, based on current contingencies in a changing environment (Mainen & Kepecs 2009; Schoenbaum & Esber 2010). These regions, thus, are less directly driven by the stimulus stream, and relate sensory information to memories and expectancies. This subtler and more indirect relationship to the stimulus stream might explain why intersubject information mapping, but not intersubject correlation mapping revealed representations shared among the subjects.

An alternative explanation concerns the spatial organization of these regions. In primary visual and auditory areas, a point-to-point correspondency between human brains can be defined with quite a high precision (1-3mm) based on retinotopic and tonotopic mapping, respectively. In higher regions, including anterior temporal and orbitofrontal cortex, there is little evidence for a comparably precise intersubject correspondency. Intersubject information mapping is robust to local misalignment of the activity patterns, relating instead the information content of the patterns within a given region contained in the searchlight. It might thus be able to reveal information shared between subjects even in the absence of a common spatial layout of the representations.

### 4.2 A method for discovering a region's stimulus-driven representational content?

The finding of intersubject information in a brain region during a common stimulus stream indicates that some aspect of the stimulus stream is represented across subjects. It does not reveal *what* aspect of the stimulus stream is represented. In order, to find out what aspect of the stimulus is represented in a region, we need to related the brain representation back to the stimulus variation over time.

This brings us back to our original consideration: We motivated analyzing intersubject information by arguing that direct analysis of the stimulus-response relationship (on which interpretations in terms of representational content would have to be based) is difficult. Natural stimulus streams are hard to model because they vary along many dimensions, and any model would have to focus on predefined dimensions, thus introducing bias. One approach is to find the dimensions of the stimulus stream that explain most variance in the response patterns (e.g. Kriegeskorte et al. 2008a, 2008b). This general idea could be applied to natural stimulus streams, for example by principal component analysis of a region's spatiotemporal pattern. The temporal mode of the first principal component could be related to the stimulus stream in an exploratory fashion. We could review the parts of the stimulus stream corresponding to high and low periods of the first-principal-component time course and attempt to infer the corresponding stimulus dimension. This would suggest the hypothesis that the stimulus dimension in question is represented in the region.

However, as discussed above high-level regions like anterior temporal and orbitofrontal cortex are believed to relate sensory information to endogenous dynamics including memories, learned reward contingencies, and current goals. We therefore expect that a substantial proportion of their response variance is not directly driven by the stimulus stream. The first-principal-component time course would, thus, be expected to strongly reflect endogenous along with stimulus-driven dynamics. Intersubject canonical correlation analysis can help us constrain the exploratory process to the stimulus-driven component of the dynamics. For example, we could use the time courses associated with the first canonical correlation (Fig. 2b), instead of the first-principal-component time course, to attempt to discover the underlying stimulus dimension commonly represented in the subjects.

In conclusion, intersubject information mapping can reveal regions representing the same stimulus information across subjects without the need for a descriptive model of the stimulus stream. It therefore constitutes a promising direction for discovering population-code representations of stimulus streams and their representational content, especially as we move toward more complex and natural stimulus streams and toward higher-resolution and higher-dimensional response measurements.

## 5 Acknowledgements

The FMRI data were collected by the Experience-based Cognition Project (Walter Schneider, U of Pittsburgh) and made publicly available in the context of the Pittsburgh Brain Activity Interpretation Competition 2006.

